# SVD-SN2N: Self-Inspired Noise2Noise Learning for Denoising Log-Compressed SVD-Filtered Ultrasound Imaging

**DOI:** 10.64898/2026.09.10.750668

**Authors:** Ge Zhang, Eric Buffle, Nicolas Zucker, Thomas Deffieux, Nathalie Ialy-Radio, Mathieu Pernot, Sophie Pezet, Mickaël Tanter

**Affiliations:** Physics for Medicine Paris, INSERM U1273, ESPCI Paris, PSL University, CNRS, 75012 Paris, France

**Keywords:** Contrast-enhanced ultrasound, log-compression, Noise2Noise, power Doppler, self-supervised learning, singular value decomposition, ultrasound denoising

## Abstract

Singular value decomposition (SVD) is the reference clutter-rejection strategy for medical ultrafast ultrasound imaging, yet SVD-filtered images retain a residual noise floor that obscures microvascular signals at depth. Supervised denoising cannot address this gap because clean references do not exist, and most physics-based alternatives require radio-frequency or in-phase/quadrature data that clinical scanners do not expose. We introduce SVD-SN2N, a self-supervised Noise2Noise framework that operates entirely on the post-SVD image and requires neither clean targets nor channel-level access. Our central contribution is to show that the residual fluctuations of SVD-filtered images are dominated by multiplicative speckle, which violates the zero-mean additive assumption of Noise2Noise theory, and that a single log-compression step converts it into an additive perturbation whose variance no longer depends on the signal, restoring the conditions the framework requires. The log-compressed image is split into two statistically independent half-size copies by diagonal 2×2 resampling, rescaled by Fourier zero-padding interpolation, and used to train a U-Net under a self-constrained twin-prediction loss. We assess the underlying assumptions directly by measuring the residual-noise distribution, its spatial correlation and the signal similarity within diagonal pixel pairs, and we validate the framework on four heterogeneous datasets spanning preclinical and clinical, contrast-enhanced and contrast-free, and 2-D to 3-D regimes. Relative to conventional SVD, SVD-SN2N raises image-derived SNR by 4.2–10.6 dB and narrows the apparent vessel full-width at half-maximum by 34%–59%, providing a post-SVD denoising front-end compatible with the image-only data available on clinical scanners.

## 1. Introduction

ULTRAFAST ultrasound imaging, based on coherent compounding of plane- or diverging-wave transmissions, has transformed microvascular imaging by enabling acquisitions at thousands of frames per second over large fields of view [1], [2]. Combined with singular value decomposition (SVD) clutter filtering [3], ultrafast acquisitions give access to slow blood-flow signals that conventional high-pass Doppler filters cannot recover, and underpin techniques such as functional ultrasound imaging, ultrasensitive power Doppler and ultrasound localization microscopy (ULM) [1], [4], [5]. SVD exploits the difference in spatiotemporal coherence between tissue and blood: after reshaping the dataset into a Casorati matrix, the first singular vectors capture slowly varying, spatially coherent tissue motion, the last singular vectors describe electronic and thermal noise, and an intermediate range carries the blood-flow signal [3], [6].

Despite this elegant separation, the boundary between the blood and noise subspaces is rarely clean, so small signals at depth remain masked by a speckle-like background even after SVD [7], [8]. A supervised network could in principle suppress this residual noise [9], [10], but supervision is ill-posed here: no pixel-accurate ground truth exists, clinical acquisitions are short so curated training sets are necessarily small, and the microvascular structure the network would have to learn is precisely the unknown of interest. The usual workaround is to synthesize training pairs with physics-based simulators such as Field II or k-Wave, in which a known scatterer distribution is propagated through an idealized acoustic model. Models trained on such pairs inherit the simplifying assumptions of that model— homogeneous speed of sound, weak scattering, idealized aberration, stationary scatterer fields—and generalize poorly to *in-vivo* data. A more robust learning target is therefore the residual noise pattern itself, which is far more stationary across acquisitions than the anatomy and is consequently better suited to self-supervised training on a small, real, *in-vivo* dataset.

A second motivation comes from clinical translation. The most effective non-learning approaches to ultrasound noise and resolution share one structural property: they exploit coherence among raw acquisition channels, either at transmission—amplitude-modulated SVD (AM-SVD) [11], [12] and nonlinear sound-sheet microscopy [13]—or at processing—xDoppler [14], frame multiply-and-sum [15], short-lag spatial coherence [16] and delay-multiply-and-sum [17]. All of them require the transmit/receive channel data or the pre-beamforming IQ stack with its phase information. Commercial clinical scanners expose neither: they deliver post-beamforming, post-compression B-mode, Doppler or contrast images, typically through a DICOM export. Any denoising tool that requires RF or channel-level data is therefore structurally unable to help the clinicians who need it most. A framework operating directly on the displayed image is, by contrast, compatible with the data clinicians actually have, requires a single short acquisition and no clean reference, and can be retrained patient-by-patient on whatever the scanner delivers.

Self-supervised denoising offers a way to avoid clean references. The Noise2Noise (N2N) result showed that a network fed with two independently corrupted realizations of the same signal converges to the same solution as one trained on clean targets, provided the noise is zero-mean and statistically independent across the two copies [18]. Variants such as Noise2Void [19], Noise2Self [20] and pipelines developed for calcium imaging [21] or fluorescence microscopy [22], [23] using U-Net [24] have shown that paired noisy frames are not even required if the noise is pixel-wise independent. In ultrasound, self-supervised denoising has been explored on RF and B-mode data [25], but to our knowledge no study has targeted the residual noise of SVD-filtered ultrafast images, nor investigated the pre-processing required to make self-supervised theory applicable to it.

A direct transposition of N2N to SVD-filtered power Doppler is in fact problematic, because the residual fluctuations of those images are dominated by speckle-like, signal-dependent (multiplicative) noise, which is incompatible with the zero-mean additive noise assumed by N2N theory [26]. A central methodological contribution of this work is to show that the mismatch is resolved by a single, well-understood pre-processing step: after log-compression the multiplicative speckle becomes an additive perturbation with a signal-independent variance, restoring the assumptions of N2N. Building on this observation we introduce SVD-SN2N, which combines (i) a log-compression step that renders the residual speckle additive with a signal-independent variance; (ii) a diagonal resampling operation that splits every 2×2 pixel block into two statistically independent, spatially matched half-size companions, followed by a band-limited Fourier zero-padding interpolation restoring the original grid; and (iii) a U-Net trained with a self-constrained loss combining two N2N cross-prediction terms with a twin-prediction consistency term. A Patch2Patch augmentation increases the effective number of training patches extracted from a single short acquisition.

Because these three ingredients rest on explicit statistical assumptions, we do not take them for granted: Section 2.8 reports direct measurements of the residual-noise distribution, of its spatial correlation, and of the signal similarity within the diagonal pixel pairs. We then validate the framework on four datasets spanning preclinical and clinical, contrast-enhanced and contrast-free, and 2-D to 3-D regimes (Table 1). On the two contrast-enhanced 2-D datasets, a three-way comparison—conventional SVD, SVD-SN2N trained on a linear-scale input, and SVD-SN2N trained on a log-compressed input—isolates the contribution of log-compression, since the linear-scale control is identical in every other respect. The gain grows systematically as the baseline SNR decreases, which is the signature of a multiplicative-noise process, while the contrast-free 2-D and 3-D row-column-addressed (RCA) datasets show that the same behaviour carries over to other acquisition geometries and contrast regimes.

**Table 1.** Summary of the Four Datasets Used in This Study.

|  | Dataset 1 | Dataset 2 | Dataset 3 | Dataset 4 |
| --- | --- | --- | --- | --- |
| Subject | Rat brain | Human breast | Rat brain | Rat brain |
| Dimensionality | 2-D | 2-D | 2-D | 3-D |
| Contrast agent | SonoVue | SonoVue | None | SonoVue |
| Imaging system | Verasonics Vantage | Mindray Resona R9 | Iconeus One | Verasonics Vantage |
| Frame rate | 1000 Hz | 80 Hz | 500 Hz | 333 Hz |
| Transmission frequency | 15.625 MHz | 6 MHz | 15 MHz | 10 MHz |

Datasets 1 and 2 are 2-D contrast-enhanced acquisitions used for both qualitative and quantitative validation. Datasets 3 and 4 extend the qualitative validation to the contrast-free 2-D regime and to volumetric 3-D imaging.

## 2. Materials and Methods

### 2.1 Dataset 1—In-Vivo Rat-Brain Ultrafast Microbubble Power Doppler

Dataset 1 is the public benchmark released by Heiles *et al*. [7]. Ultrafast acquisitions of the rat brain were performed through a cranial window after bolus injection of microbubbles, with coherent plane-wave compounding at 5 angles from −6° to +6°; complex IQ stacks were made publicly available. Full details of animal preparation, acquisition and beamforming are given in the original publication. A one-second acquisition block was used to train SVD-SN2N and an independent four-second block of the same session was used to test it.

### 2.2 Dataset 2—Clinical Human Breast CEUS

Clinical contrast-enhanced ultrasound (CEUS) acquisitions of human breast tumors were obtained in previous studies [27], [28] on a Mindray Resona R9 system (Mindray, Shenzhen, China) with an L11-3U linear probe, by a physician with more than twenty years of breast-ultrasound experience. The patient was supine with the ipsilateral arm raised. Contrast imaging was performed at a mechanical index of 0.08 after a bolus of SonoVue microbubbles (Bracco, Milan, Italy) injected via the cubital peripheral vein and flushed with saline. Frames were recorded during microbubble wash-in with a real-time dual display of B-mode and contrast images used to keep the imaging plane fixed. Critically, the contrast images were exported from the clinical scanner directly in DICOM format: only the device-rendered, post-beamforming image stack was available, not the underlying RF or IQ data, so SVD-SN2N is evaluated here on exactly the kind of data a clinician obtains in routine practice. A 2 s block of consecutive DICOM frames was used for training and an independent 10 s block of the same session for testing.

### 2.3 Dataset 3—In-Vivo Rat-Brain Contrast-Free Ultrafast Power Doppler

*In-vivo* contrast-free ultrafast power Doppler of the rat brain was acquired with a 1-D linear transducer (128 elements, 110 µm pitch) connected to a functional ultrasound scanner (Iconeus One, Iconeus, Paris, France, in collaboration with INSERM ART Biomedical Ultrasound, Paris, France). Each ensemble consisted of 11 plane waves tilted from −10° to +10° at a 5.5 kHz pulse-repetition frequency, coherently compounded into 2-D images [29], [30]. SVD-SN2N was trained on a one-second block and tested on independent acquisitions from the same animal. This dataset extends the validation to the contrast-free regime, where the residual noise of the SVD output is dominated by speckle and electronic noise rather than by sparse microbubble events.

### 2.4 Dataset 4—In-Vivo 3-D Rat-Brain Harmonic Imaging Using a Row-Column-Addressed Array

*In-vivo* 3-D contrast-enhanced harmonic imaging of the rat brain was performed on a research scanner (Verasonics Vantage, Verasonics, USA) driving a row-column-addressed (RCA) array (160 elements; 15.625 MHz central frequency; 80% bandwidth at −6 dB), with acquisition and beamforming code developed in MATLAB (MathWorks, USA). The HAM-SVD sequence transmitted three half-cycle plane waves with 7 angles each on the row and column apertures, spanning −2.25° to +2.25°. Transmissions were repeated for four duty-cycle levels (DC_1_ = 0.55, DC_2_ = 0.70, DC_3_ = 0.85, DC_4_ = 1.00), the duty cycle being the ratio of pulse duration to pulse-repetition period; the mechanical index is 0.025 for DC_4_. Higher duty cycles transfer more acoustic energy and effectively modulate the transmit amplitude, giving nonlinear responses at four pressure levels. Channel data were beamformed offline on GPU with a delay-and-sum scheme on a λ/2 grid. SVD-SN2N was applied frame-by-frame on the three orthogonal cross-sections extracted from the volumetric power-Doppler maps, with no modification other than the slice-by-slice application. Table 1 summarizes the four datasets.

### 2.5 Conventional SVD Clutter Filtering

For each ensemble of *N* slow-time frames, the data were reshaped as a Casorati matrix *C* of size (*N*_x_·*N*_z_) × *N*, where *N*_x_ and *N*_z_ are the lateral and axial pixel counts. For Datasets 1, 3 and 4 the matrix was built from the complex IQ stack; for the clinical breast acquisition the IQ stream was not exported, so the matrix was built directly from consecutive DICOM contrast frames, matching the data path available to any user of a commercial system. After computing *C* = *USV*\*, the tissue subspace was suppressed by discarding the first singular modes up to a threshold *T*_1_ taken at the inflection point of the normalized singular-value energy curve, the number of frames serving as the second threshold *T*_2_. The blood-flow image was reconstructed from the retained modes and its amplitudes integrated over the ensemble to produce a power-Doppler map. For the breast CEUS acquisition, an additional temporal incoherent summation across consecutive ensembles built the microbubble-weighted map used as input to SVD-SN2N.

### 2.6 Log-Compression of SVD-Filtered Images

The residual fluctuations of SVD-filtered power-Doppler images are dominated by speckle-like, signal-dependent noise. For fully developed speckle the envelope follows a multiplicative model

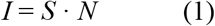

where *S* is the blood-signal amplitude, *N* a unit-mean, Rayleigh-distributed speckle factor and *I* the observed pixel value [26]. The noise variance therefore scales with the signal, so neither the zero-mean nor the additive assumption of the N2N theorem [18] holds in the linear-scale domain. Taking the logarithm decouples the two terms,

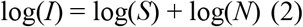

and log(*N*) satisfies the conditions the N2N objective requires: (i) its mean does not depend on the local signal *S* and is absorbed by a global offset, so it is zero-median and zero-mean after de-biasing; (ii) for Rayleigh-distributed *N* it follows a Fisher–Tippett (log-Rayleigh) law whose variance, π^2^/24 ≈ 0.411 in natural-log units, no longer depends on the local signal; and (iii) the logarithm being a pointwise operation, it leaves the spatial correlation structure of *N* unchanged. What log-compression buys is therefore not Gaussianity but additivity with a signal-independent variance, and this is what the objective needs: the loss of (3) is an *ℓ*_1_ loss and converges to the conditional median of its target, so a zero-median residual that is independent across the two copies suffices and normality is not required. Conditions (i)–(iii) are measured on our own data in Section 2.8. In practice we apply a 20·log_10_(·) operation followed by clipping to a prescribed dynamic range (typically 50–100 dB) and min/max normalization to [0, 1]; the remainder of the pipeline is identical for linear- and log-scale inputs, so the three-way comparisons of Section 3 differ only in this pre-processing choice.

### 2.7 SVD-SN2N Framework

SVD-SN2N takes as input a single log-compressed SVD-filtered power-Doppler or contrast-weighted image *y* and outputs a denoised estimate 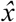 of the underlying log-signal. It rests on three design pillars: self-supervised generation of two statistically independent noisy copies from the single input; a U-Net trained with a self-constrained N2N loss; and a Patch2Patch augmentation tailored to ultrasound time series. The pipeline is summarized in Fig. 1.

**Fig. 1.**
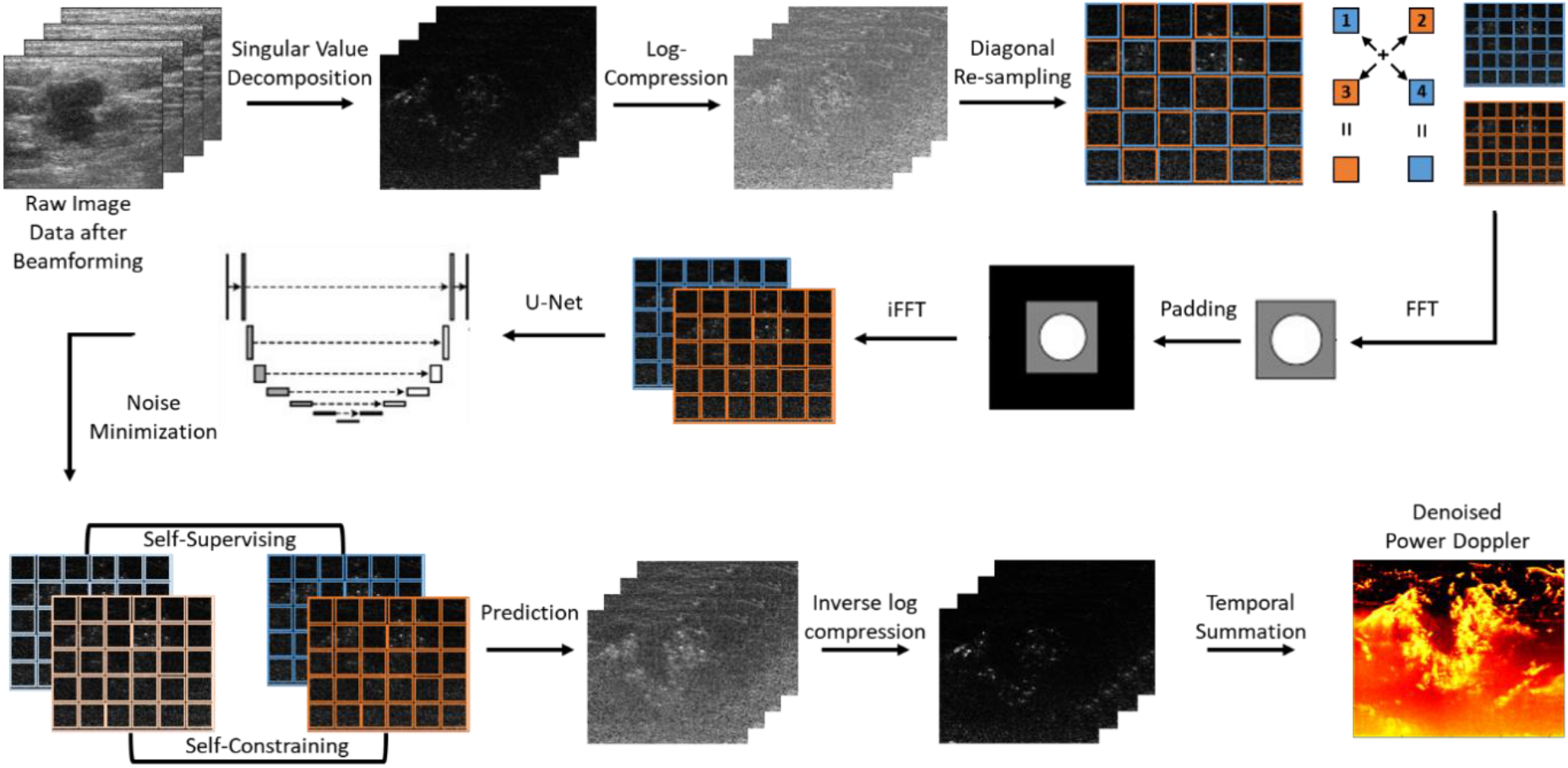
Overview of the SVD-SN2N processing pipeline. Raw ultrafast IQ data are passed through a conventional SVD clutter filter to remove tissue signal and isolate blood-flow components (top-left). The filtered image is log-compressed, augmented by a Patch2Patch scheme, and split into two statistically independent half-size copies by diagonal resampling of every 2×2 pixel block. A Fourier zero-padding interpolation rescales each copy back to the original grid, so that the paired images share the same structural scale as the input. The resulting twin noisy images are fed to a U-Net trained with a self-constrained N2N loss enforcing both cross-prediction agreement (self-supervising) and twin-prediction consistency (self-constraining).

#### Noise Model

After log-compression we adopt the additive observation model *y* = *x* + *n*, where *x* = log(*S*) is the unknown clean log-signal and *n* = log(*N*) is zero-median after de-biasing, of signal-independent variance and approximately independent from pixel to pixel. Raw ultrasound channel noise satisfies none of this—RF acquisition, IQ demodulation and envelope detection make its statistics both signal-dependent and non-stationary—but the combination of SVD clutter filtering and log-compression places the residual in the regime the self-supervised objective requires. This is why SVD-SN2N is designed around log-compressed image-level data rather than RF or IQ signals.

#### Diagonal Resampling

We split the log-compressed image into two half-size companions sharing the same underlying vascular structure but differing in their noise realization. Ultrafast Doppler images are sampled on a grid finer than half the diffraction limit, so every local point spread function (PSF) occupies at least a 2×2 pixel block. Denoting by *p*_1_, *p*_2_ (first row) and *p*_3_, *p*_4_ (second row) the four pixels of such a block, we form the diagonal averages *x*_1_ = (*p*_1_ + *p*_4_)/2 and *x*_2_ = (*p*_2_ + *p*_3_)/2 and scan the operation across the image to produce two half-resolution subimages. Under the additive-independent model, the residual noises of *x*_1_ and *x*_2_ are built from the disjoint pixel sets [*p*_1_, *p*_4_] and [*p*_2_, *p*_3_], so Cov[*n*_1_, *n*_2_] vanishes exactly when the residual is white; in practice it does not vanish, and Section 2.8 quantifies what remains. The diagonal pairing is nevertheless the natural choice, for a reason that is exact: among the three two-way partitions of a 2×2 block, it is the only one whose two companions share the same centroid, so it introduces no half-pixel shift and the underlying signal is sampled at the same location in *x*_1_ and *x*_2_. Compared with the temporal resampling proposed for calcium or voltage imaging [31], which requires two consecutive frames of an identical underlying signal, diagonal resampling operates inside a single image and therefore remains valid for fast bubble dynamics, respiratory motion and arbitrary ensemble lengths.

#### Fourier Zero-Padding Interpolation

Because *x*_1_ and *x*_2_ are twice smaller than the input, they are rescaled back to the original grid before training. Spatial-domain interpolation such as bilinear or bicubic upsampling would reconstruct missing pixels as linear combinations of noisy neighbors and thereby break the local independence of the noise. We instead use Fourier zero-padding interpolation: each subimage is mirror-padded to prevent boundary artifacts, Fourier-transformed, zero-padded symmetrically in the frequency domain so that its spectrum occupies twice as many samples in each direction without exceeding the original transfer-function support, inverse-transformed and cropped back to the original size. This is an exact band-limited interpolation that respects the transfer function of the imaging system and introduces no pixel-to-pixel correlation.

#### Self-Constrained N2N Loss

Given *x*_1_ and *x*_2_, the U-Net *f*_θ_ is trained with

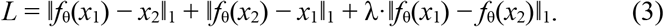

The first two terms are the standard N2N cross-prediction losses and push each prediction toward the complementary noisy target. The third is a self-consistency regularizer: because *x*_1_ and *x*_2_ describe the same underlying vascular signal, their denoised versions should coincide, and forcing agreement empirically improves denoising quality when only a single training frame is available. The weight λ is set to 1 by default. Note that the loss target is the companion noisy pixel, not a clean reference: the network is driven to learn the pixel-wise noise pattern, which is shared in distribution between the two copies, rather than the vascular signal itself.

#### Patch2Patch Data Augmentation and Train/Test Separation

Training patches of 128 × 128 pixels are drawn *exclusively* from the training block of each dataset. The independent test block contributes no patch at any stage, is never used for model selection or for early stopping, and is processed only once, after training has ended; training and test blocks are disjoint in time within the same acquisition session. Augmentation consists of three patch-exchange operations, drawn at random: (i) exchange of two patches at the same spatial location between two different time points of the training block; (ii) exchange of two patches at two different spatial locations within the same frame; and (iii) exchange of two patches between two training blocks acquired from the same subject with the same probe, transmit sequence and post-processing settings, so that the residual-noise statistics of the two sources are identical. Blocks differing in any of these acquisition parameters are never mixed, since a change of noise distribution would invalidate the N2N assumption. Each patch is additionally subjected to a random multiple of 90° rotation or to a horizontal or vertical flip. All three operations permute patches without modifying pixel values, so the intrinsic noise distribution of the data is preserved.

### 2.8 Empirical Assessment of the Noise Assumptions

The self-supervised objective rests on three assumptions that we assess directly rather than presume: that the residual *n* of the log-compressed SVD output is zero-median and of signal-independent variance; that it is spatially independent at the pixel scale; and that the underlying signal is identical in the two diagonal companions. Three diagnostics were computed on vessel-free and high-SNR regions of the contrast-enhanced datasets.

#### (i) Residual distribution

Within a vessel-free background region, the residual was obtained by subtracting a local median estimate of log(*S*). Its histogram and a quantile–quantile plot against a Fisher–Tippett law predicted for L-look speckle, together with a Gaussian of matched variance, identify which law it follows, and the measured variance is compared with the value π^2^/24 ≈ 0.41 predicted for fully developed Rayleigh speckle. *(ii) Spatial independence*. The 2-D normalized autocorrelation of the same residual was computed, and its width at 1/*e* quantifies the residual correlation length in pixels; a length below one pixel supports the pixel-wise independence assumption, whereas a larger value indicates partially correlated speckle. *(iii) Signal similarity within diagonal pairs*. On high-SNR regions we computed the Pearson correlation between *x*_1_ and *x*_2_ and the distribution of their difference; an identical underlying signal with independent noise implies a correlation close to unity and a difference centred on zero. Results are reported in Section 3.1.

### 2.9 Implementation and Inference

The network *f*_θ_ is a 2-D U-Net with four down-sampling and four up-sampling stages, 3×3 convolutions, batch normalization, Leaky ReLU activations, symmetric skip connections and a 1×1 sigmoid output head. Training was performed on a single NVIDIA GeForce RTX 3080 Ti GPU under Python 3.7.6 / PyTorch 1.7.1 / CUDA 11.6. Hyperparameters are summarized in Table 2.

**Table 2.** Summary of SVD-SN2N Training Hyperparameters.

| Hyperparameter | Value |
| --- | --- |
| Input pre-processing | SVD clutter filter $\rightarrow 20 \cdot \log_{10}(\cdot)$ (log runs only) |
| Input patch size | $128 \times 128$ pixels |
| Batch size | Adaptive (GPU-memory limited) |
| Optimizer | Adam ($\beta_1, \beta_2$) = (0.5, 0.999) |
| Learning rate | $2 \times 10^{-4}$ |
| Self-consistency weight $\lambda$ | 1 |
| Epochs / iterations | $\approx 100 / 4000$ |
| Hardware / framework | NVIDIA RTX 3080 Ti; Python 3.7.6 / PyTorch 1.7.1 / CUDA 11.6 |

At inference, two predictions *f*_θ_(*x*_1_) and *f*_θ_(*x*_2_) are produced from the two diagonal-resampling copies of the same input, and their mean is used as the final denoised image. Their pixel-wise absolute difference σ(*r*) = |*f*_θ_(*x*_1_)(*r*) − *f*_θ_(*x*_2_)(*r*)| is available at no additional cost and is reported here as a *self-consistency map*: it measures how much the network disagrees with itself, typically on faint or ambiguous vessels. It is a relative, uncalibrated indicator and is not a validated estimate of predictive uncertainty; calibrating it against a known reference is left to future work.

### 2.10 Three-Way Experimental Design and Quantitative Metrics

For each contrast-enhanced 2-D dataset (Datasets 1 and 2), three conditions were compared: conventional SVD, SVD-SN2N trained on a linear-scale input, and SVD-SN2N trained on a log-compressed input. The linear-scale condition is a control isolating the contribution of log-compression: architecture, loss, augmentation and optimizer were identical across conditions and only the input pre-processing differed. Two image-derived metrics were used. Signal-to-noise ratio (SNR, dB) was computed as SNR = 20·log_10_(μ_v_/σ_b_) on manually drawn vessel (ROI v) and vessel-free background (ROI b) regions shared between the three methods (Table 3). Vessel full-width at half-maximum (FWHM) was obtained by fitting a 1-D Gaussian to every lateral and axial profile crossing isolated vessels, over a large number of automatically extracted line profiles, and summarized by mean, median and standard deviation (Table 4); the same set of lines was used for all three methods. Both metrics are computed on the images themselves and therefore quantify apparent contrast and apparent vessel width; neither establishes that the recovered structures are faithful to the underlying microvasculature, a distinction we return to in Section 4. For Datasets 3 and 4 the comparison is restricted to qualitative side-by-side maps and intensity profiles, since these acquisitions do not provide a comparable set of paired vessel/background ROIs across all conditions.

**Table 3.** SNR (dB) at Two Vessel/Background ROI Pairs per Dataset.

| ROI | Conventional<br>SVD | SVD-SN2N<br>(linear) | SVD-SN2N<br>(log) |
| --- | --- | --- | --- |
| Rat brain—ROI 1 | 20.13 | 22.56 | 24.32 |
| Rat brain—ROI 2 | 13.74 | 15.52 | 23.89 |
| Breast—ROI 1 | 12.54 | 14.23 | 21.34 |
| Breast—ROI 2 | 5.87 | 6.98 | 16.47 |

**Table 4.** Vessel FWHM Distributions (mm)

| Method | Mean | Median | Std. dev. |
| --- | --- | --- | --- |
| *Rat brain—axial* |  |  |  |
| Conventional SVD | 0.531 | 0.354 | 0.448 |
| SVD-SN2N (linear) | 0.607 | 0.358 | 0.533 |
| SVD-SN2N (log) | 0.312 | 0.207 | 0.319 |
| *Rat brain—lateral* |  |  |  |
| Conventional SVD | 0.754 | 0.596 | 0.469 |
| SVD-SN2N (linear) | 0.710 | 0.556 | 0.463 |
| SVD-SN2N (log) | 0.309 | 0.293 | 0.241 |
| *Breast—axial* |  |  |  |
| Conventional SVD | 0.793 | 0.531 | 0.622 |
| SVD-SN2N (linear) | 0.776 | 0.490 | 0.623 |
| SVD-SN2N (log) | 0.527 | 0.303 | 0.492 |
| *Breast—lateral* |  |  |  |
| Conventional SVD | 0.859 | 0.623 | 0.613 |
| SVD-SN2N (linear) | 0.792 | 0.506 | 0.610 |
| SVD-SN2N (log) | 0.454 | 0.270 | 0.467 |

## 3 Results

### 3.1 Validity of the Noise Assumptions and the Role of Log-Compression

The practical effect of log-compression is shown in Fig. 2(a) on a representative SVD-filtered frame of the rat-brain dataset, which displays the network input, its output, and the normalized input-to-output difference—the noise pattern actually removed—when training is performed on the linear-scale signal (top row) or on the log-compressed signal (bottom row). On the linear scale the network amplifies both the bright scatterers and the surrounding speckle, leaving the output as noisy as the input, and the difference map is essentially black: the network has learned little more than an identity mapping. On the log scale the scatterers are preserved while the background noise floor drops, and the difference map displays a structured speckle pattern—the noise the network has isolated and removed.The three assumptions on which this behaviour rests were then measured directly (Section 2.8).

**Fig. 2.**
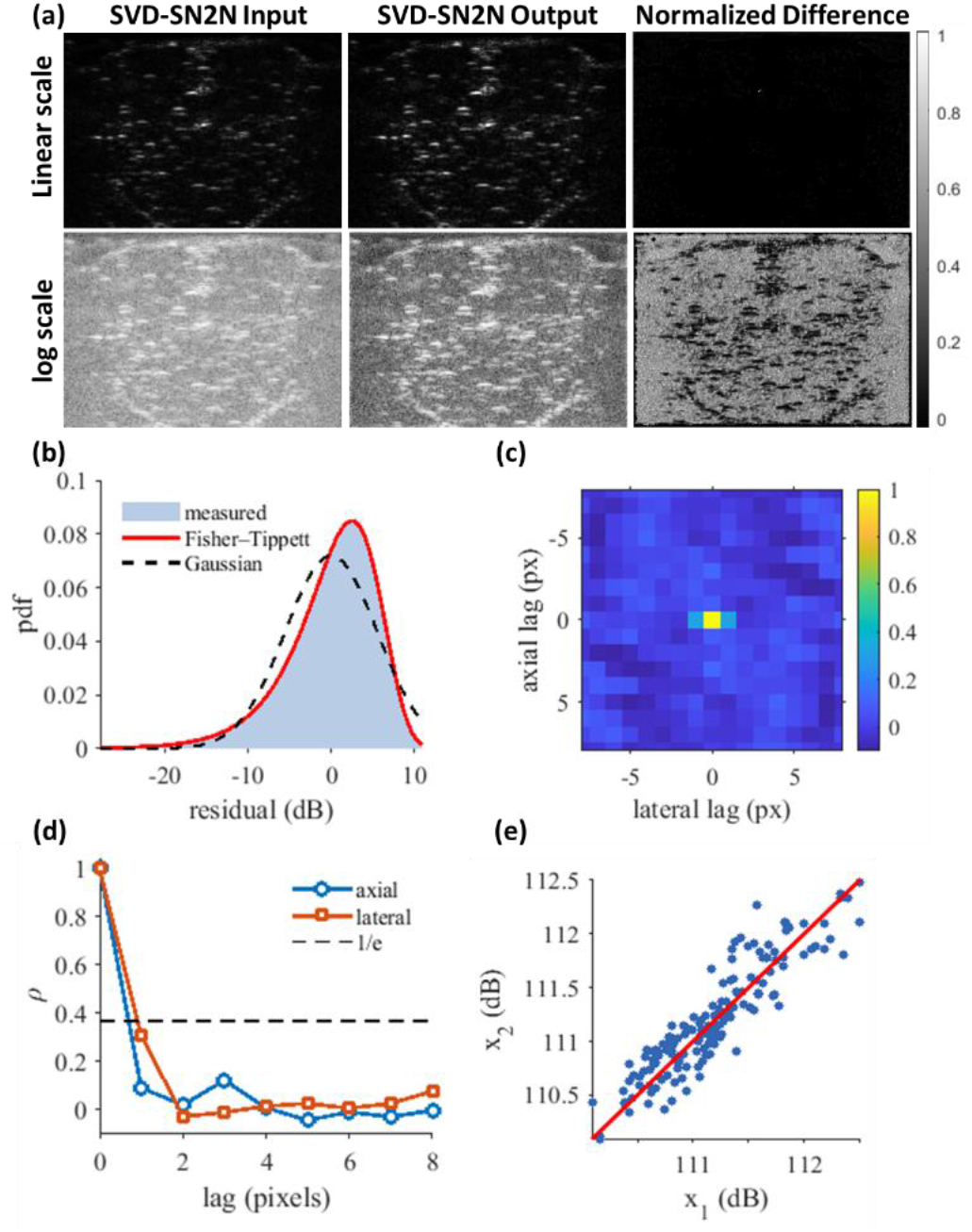
Effect of the log-compression step, and direct validation of the assumptions it relies on, on the *in-vivo* rat brain. (a) SVD-SN2N input, output and normalized input-to-output difference under linear-scale training (top row) and log-compressed training (bottom row); linear-scale training amplifies both signal and noise and yields a nearly empty difference map, whereas log-compressed training preserves the signal, lowers the background floor and yields a structured difference map that reveals the speckle actually suppressed. (b) Distribution of the log-compressed residual in a vessel-free region (651 000 pixels pooled over 1000 single-look realizations), with the Fisher–Tippett law predicted for single-look speckle (solid) and a Gaussian of matched variance (dashed). (c), (d) Two-dimensional autocorrelation of the same residual and its axial and lateral profiles, with the 1/*e* level. (e) Diagonal companions *x*_1_ and *x*_2_ on the network input over a high-SNR region, with the identity line. Numerical values are given in Section 3.1.

#### Residual distribution

Pooled over 1000 single-look realizations of a vessel-free background region (651 000 pixels), the residual of the log-compressed SVD output had a standard deviation of 5.56 dB, that is a variance of 0.409 in natural-log amplitude units, against the 0.411 predicted by π^2^/24 for fully developed Rayleigh speckle; the corresponding equivalent number of looks is 1.00. Log-compression therefore delivers what the framework needs: an additive residual whose variance no longer depends on the local signal. It does not deliver a Gaussian residual, and the measurement makes the distinction explicit. The skewness (−1.14) and kurtosis (5.41) of the residual match the Fisher–Tippett values expected for log-compressed single-look speckle (−1.14 and 5.39) rather than the Gaussian values (0 and 3), and a quantile–quantile fit gives *R*^2^ = 0.99999 against the Fisher–Tippett law versus 0.940 against a Gaussian [Fig. 2(b)]. As set out in Section 2.6, this does not affect the validity of the objective, since the *ℓ*_1_ loss of (3) converges to the conditional median and requires a zero-median, not a normal, residual.

#### Spatial independence

The two-dimensional autocorrelation of the same residual [Fig. 2(c) and (d)] falls to 1/*e* within 0.70 pixels axially and 0.91 pixels laterally. Nearest-neighbour correlations are 0.09 axially and 0.31 laterally, and the two lags entering the diagonal split are 0.06 and −0.01. Propagating these values through the diagonal averaging gives a residual correlation of 0.39 between the noise of *x*_1_ and that of *x*_2_. The residual is therefore close to, but not exactly, pixel-wise independent. The consequence is bounded and works in the safe direction: a correlation between the two copies means that part of the noise remains predictable from the companion and is consequently retained rather than removed, so the departure from the assumption biases the framework towards under-denoising, and cannot by itself generate structure that is not present in the input.

#### Signal similarity within the diagonal pairs

Measured on the network input over a high-SNR region [Fig. 2(e)], *x*_1_ and *x*_2_ have a Pearson correlation of 0.960, and their difference is centred on −0.02 dB with a standard deviation of 0.128 dB against the 0.143 dB expected from the propagated noise alone. The residual signal-mismatch variance is −0.004 dB^2^, i.e. below the resolution of the measurement, so the underlying signal can be treated as identical in the two companions. Taken together, the three diagnostics support the additivity and signal-similarity assumptions quantitatively, and replace the independence assumption by a measured correlation whose effect on the result is characterized rather than assumed.

### 3.2 Robustness to the Number of Training Frames

SVD-SN2N can be trained from as little as a single noisy frame. Fig. 3 shows the stability of the method with respect to training-set size on both contrast-enhanced 2-D datasets, using 1, 10, 100 and 1000 frames drawn from the respective training blocks. On the linear scale the pipeline fails to converge to a clean mapping at any training-set size; on the log scale the denoised map is already essentially at its final quality with a single training frame and remains visually stable as the training set grows. This insensitivity follows from the self-consistency term of the loss, which enforces agreement between the two twin predictions even when one frame is available, and is what enables the short training regime used throughout Sections 3.3 and 3.4.

**Fig. 3.**
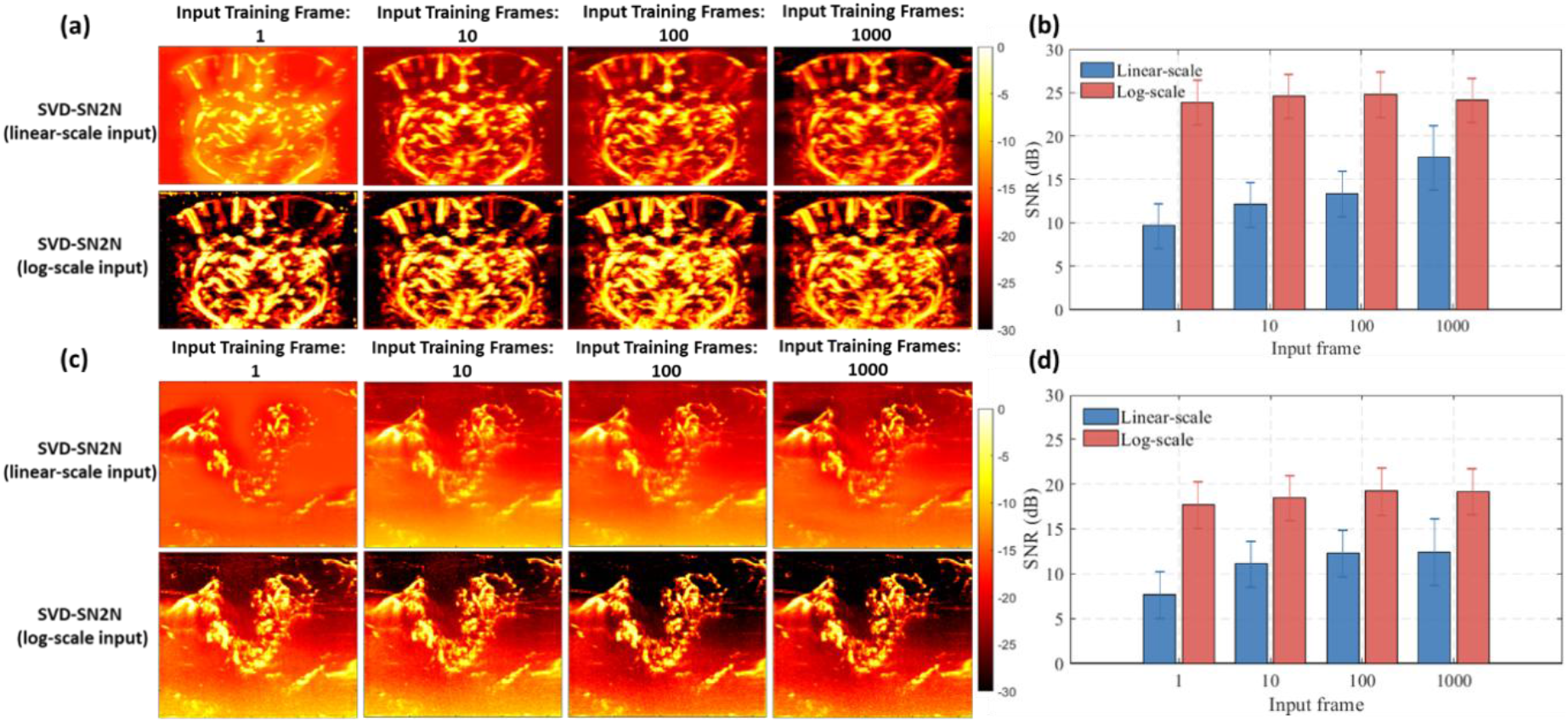
Effect of the number of training frames on SVD-SN2N for (a), (b) the rat brain and (c), (d) the human breast dataset. Training frames were drawn from the training block (1 frame in the first column, then 10, 100 and 1000 frames). For each dataset, the upper row shows SVD-SN2N trained on linear-scale inputs and the lower row shows SVD-SN2N trained on log-compressed inputs. The log-scale pipeline is already convergent from a single training frame, whereas the linear-scale pipeline fails to reach a clean mapping at any training-set size.

### 3.3 In-Vivo Rat-Brain Microbubble Power Doppler (Dataset 1)

We applied the full pipeline to the rat-brain ULM benchmark [7], training on one second of acquisition and testing on an independent four-second block. Fig. 4 compares, on the same coronal slice, conventional SVD, SVD-SN2N with linear-scale input and SVD-SN2N with log-compressed input. The linear-scale output is visually very similar to the SVD input, confirming that the multiplicative-noise regime prevents the network from learning a meaningful mapping. The log-compressed output sharpens small vessels and suppresses the speckle background, exposing cortical and subcortical microvasculature that is barely distinguishable on the conventional SVD map. On the independent test block, cortical penetrating arterioles and thalamic perforators appear more continuous, the background between vessels is strongly reduced, and deep vessels near the skull base emerge from the noise floor. The intensity profiles of Fig. 4(c) and (d) quantify this behaviour: the log-scale curve lies several decibels below the SVD and linear-SN2N curves in vessel-free portions of the profile while matching them on the vessel peaks. The same pattern recurs on every dataset below and is not described again.

**Fig. 4.**
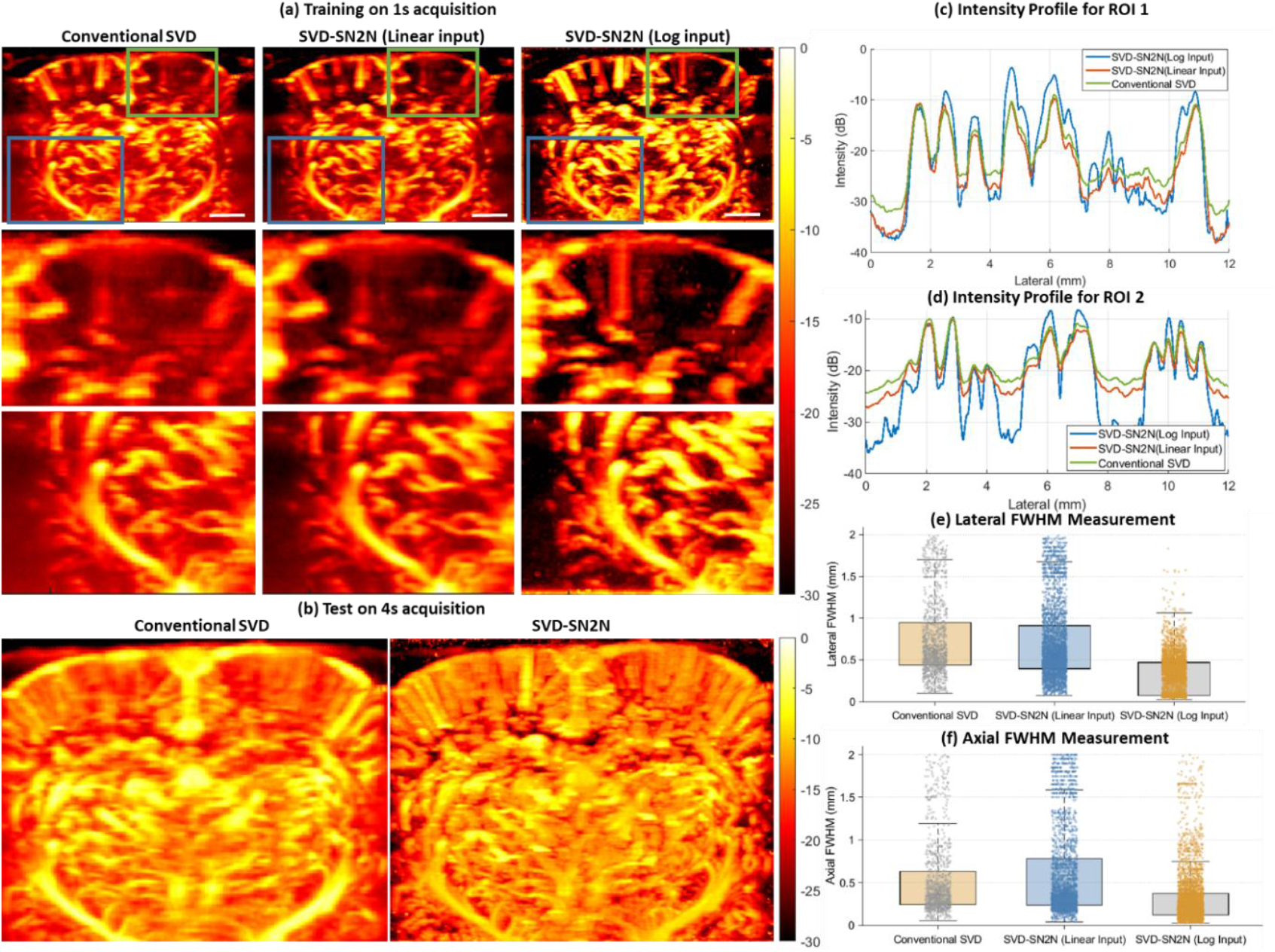
*In-vivo* rat-brain microbubble power-Doppler imaging (Dataset 1). (a) Three-way comparison on the 1 s training block: conventional SVD (left), SVD-SN2N with linear-scale input (middle), SVD-SN2N with log-compressed input (right). Top row: full coronal field of view with two regions of interest (green box: cortical region; blue box: deep thalamic region). Middle and bottom rows: zoomed views of each region. (b) Final side-by-side on the independent 4 s test block. (c), (d) Intensity profiles along lateral lines within ROI 1 and ROI 2 for the three methods. (e), (f) Lateral and axial FWHM distributions from Gaussian fits to automatically extracted vessel profiles (Table 4). All images are displayed over a 30 dB dynamic range.

### 3.4 Clinical Human Breast CEUS (Dataset 2)

We next assessed SVD-SN2N on clinical CEUS data acquired in a breast tumor patient after SonoVue injection. Here the microvascular signal is carried by sparse, rapidly moving microbubble events, a stricter test of the framework. Fig. 5 shows the three-way comparison on a representative frame, with zoom-ins on the hypervascular tumor rim and on a deep tissue region, and the test on the independent block. As in the rat brain, the linear-scale pipeline preserves the residual speckle and background leakage, whereas the log-compressed pipeline produces a cleaner map with sharper microvascular structures along the tumor rim and a substantially reduced background at depth.

**Fig. 5.**
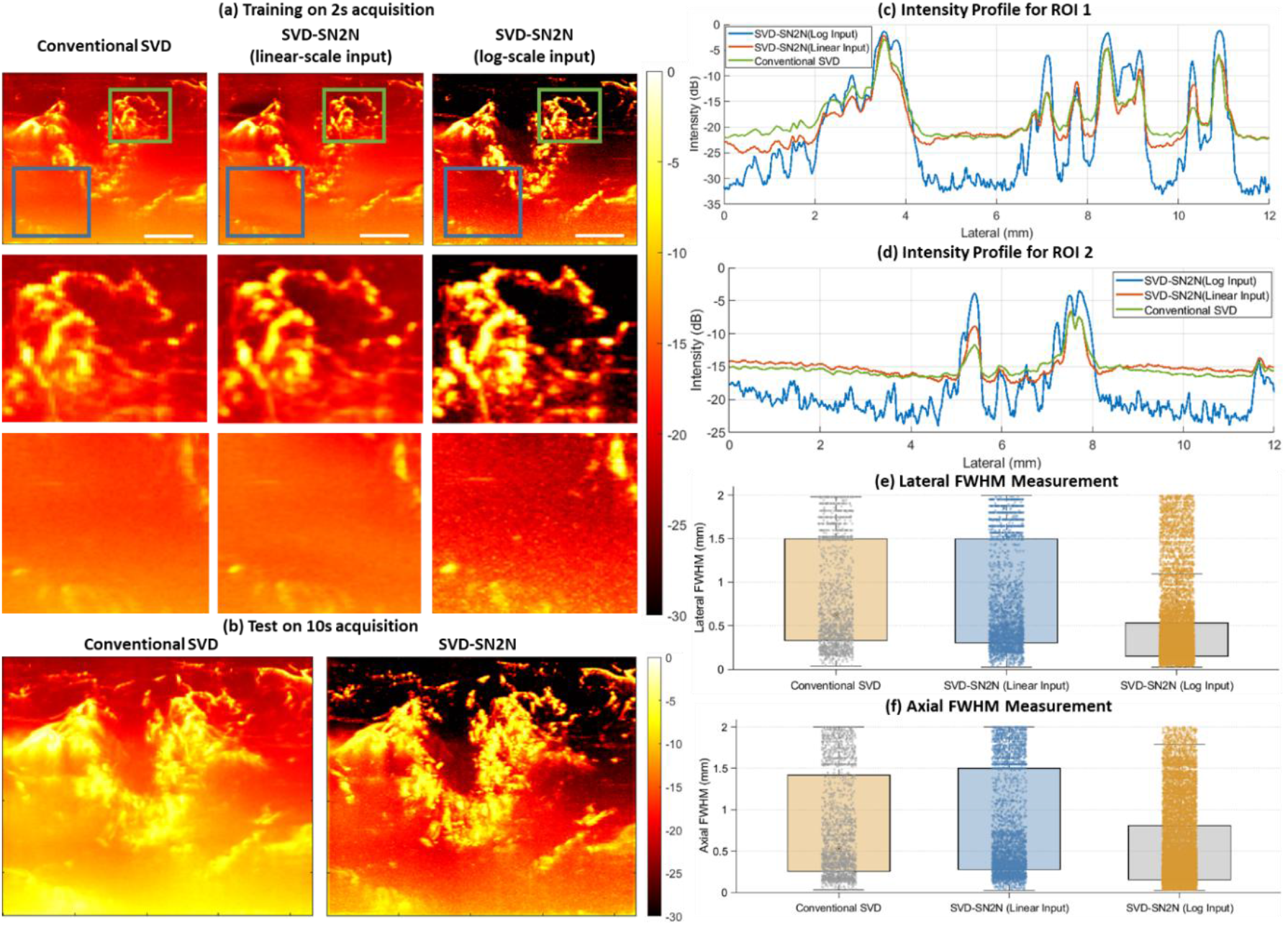
Human breast tumor CEUS imaging with SonoVue microbubbles (Dataset 2). (a) Three-way comparison on the training block: conventional SVD (left), SVD-SN2N with linear-scale input (middle), SVD-SN2N with log-compressed input (right). Top row: full field of view with two regions of interest (green box: hypervascular tumor rim; blue box: deep tissue). Middle and bottom rows: zoomed views of each region. (b) Final side-by-side on the independent test block. (c), (d) Intensity profiles along lateral lines within ROI 1 and ROI 2 for the three methods. (e), (f) Lateral and axial FWHM distributions from Gaussian fits to automatically extracted vessel profiles (Table 4). All images are displayed over a 30 dB dynamic range.

Quantitatively (Table 3), log-compressed SVD-SN2N raises the image-derived SNR by 8.80 dB in the superficial ROI 1 and by 10.60 dB in the deep ROI 2, where the conventional-SVD baseline is lowest, while the linear-scale control contributes 1.69 dB and 1.11 dB respectively. The superficial-to-deep gap accordingly narrows from 6.7 dB to 4.9 dB, so that vessels essentially buried in the noise floor at depth become directly readable. The apparent vessel FWHM (Table 4) narrows by 34% axially and 47% laterally relative to conventional SVD, with the distribution standard deviation dropping by 21% and 24%, whereas the linear-scale control leaves it essentially unchanged. These values are of the same order as those measured on the rat brain despite a very different acoustic regime, a much sparser microbubble signal and a clinical scanner exposing only DICOM contrast images.

### 3.5 In-Vivo Contrast-Free Rat-Brain Power Doppler (Dataset 3)

We then applied SVD-SN2N to contrast-free power Doppler of the rat brain (Section 2.3). No microbubbles are present and the residual SVD signal is dominated by speckle and electronic noise rather than by sparse contrast events, which makes the test stricter than for the contrast-enhanced datasets. Fig. 6 compares conventional SVD with log-compressed SVD-SN2N in power Doppler [Fig. 6(a)] and colour Doppler [Fig. 6(b)]. The log-compressed pipeline again suppresses the speckle-like background, exposing cortical microvasculature in the near field and small penetrating arterioles in the deeper thalamic region, and the intensity profiles of Fig. 6(c) reproduce the pattern described in Section 3.3. That the same behaviour persists without contrast agent indicates that the log-compression argument does not depend on the presence of microbubbles.

**Fig. 6.**
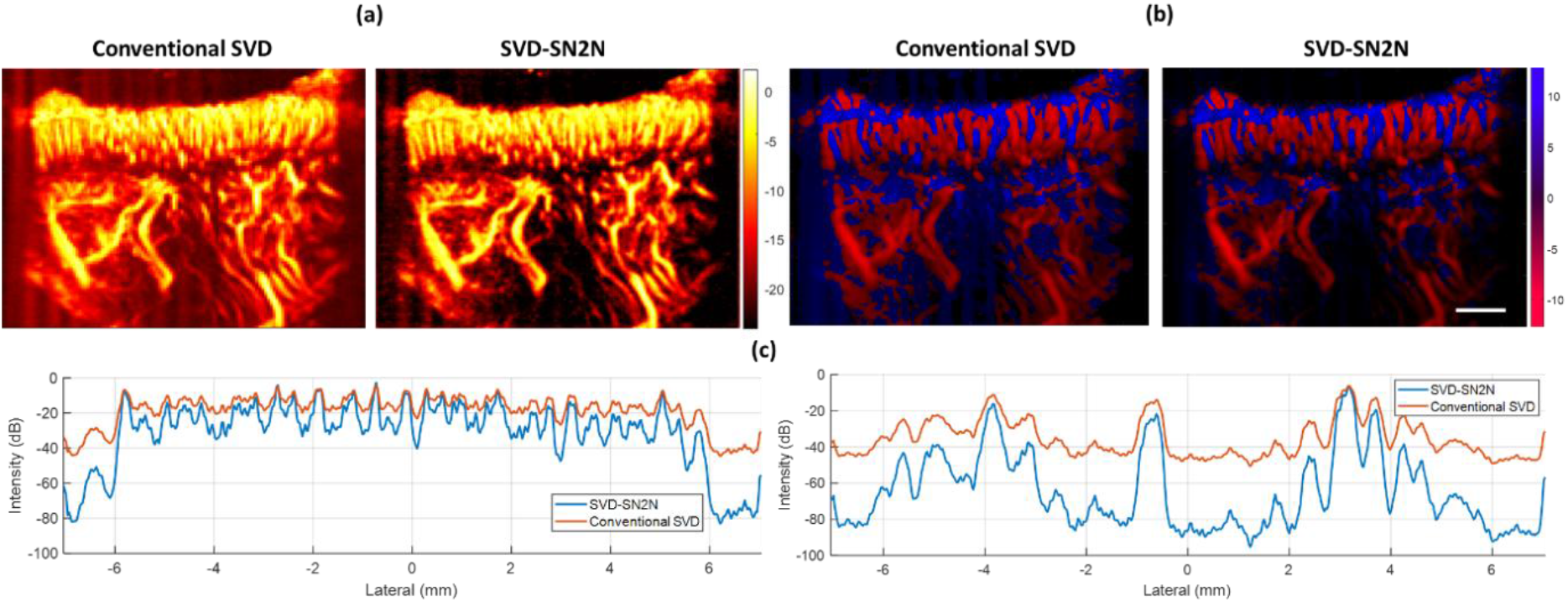
*In-vivo* contrast-free rat-brain power-Doppler imaging (Dataset 3). (a) Power Doppler and (b) colour Doppler, comparing conventional SVD with SVD-SN2N on a log-compressed input. (c) Intensity profiles along lateral lines through the near-field cortex region and in the deeper field of view. All images are displayed over a 20 dB dynamic range.

### 3.6 In-Vivo 3-D Row-Column-Addressed Harmonic Imaging (Dataset 4)

Finally, SVD-SN2N was applied to 3-D contrast-enhanced harmonic imaging of the rat brain acquired with an RCA array (Section 2.4). The volumetric power-Doppler maps were sliced into sagittal, coronal and transverse cross-sections and the framework was applied frame-by-frame on each set of slices, with no modification other than the slice-by-slice application. Fig. 7 compares conventional SVD with log-compressed SVD-SN2N over an acquisition of 50 volumes. In all three planes the speckle-like background is reduced while small vascular structures along the cortical surface and in the deeper parenchyma are preserved or sharpened, and the intensity profiles of Fig. 7(b) confirm that the decibel-level reduction of the inter-vessel background observed in 2-D translates directly to 3-D. The framework is therefore not tied to a particular probe geometry and extends to volumetric imaging, including with the cross-shaped PSF and lower base SNR characteristic of RCA arrays.

**Fig. 7.**
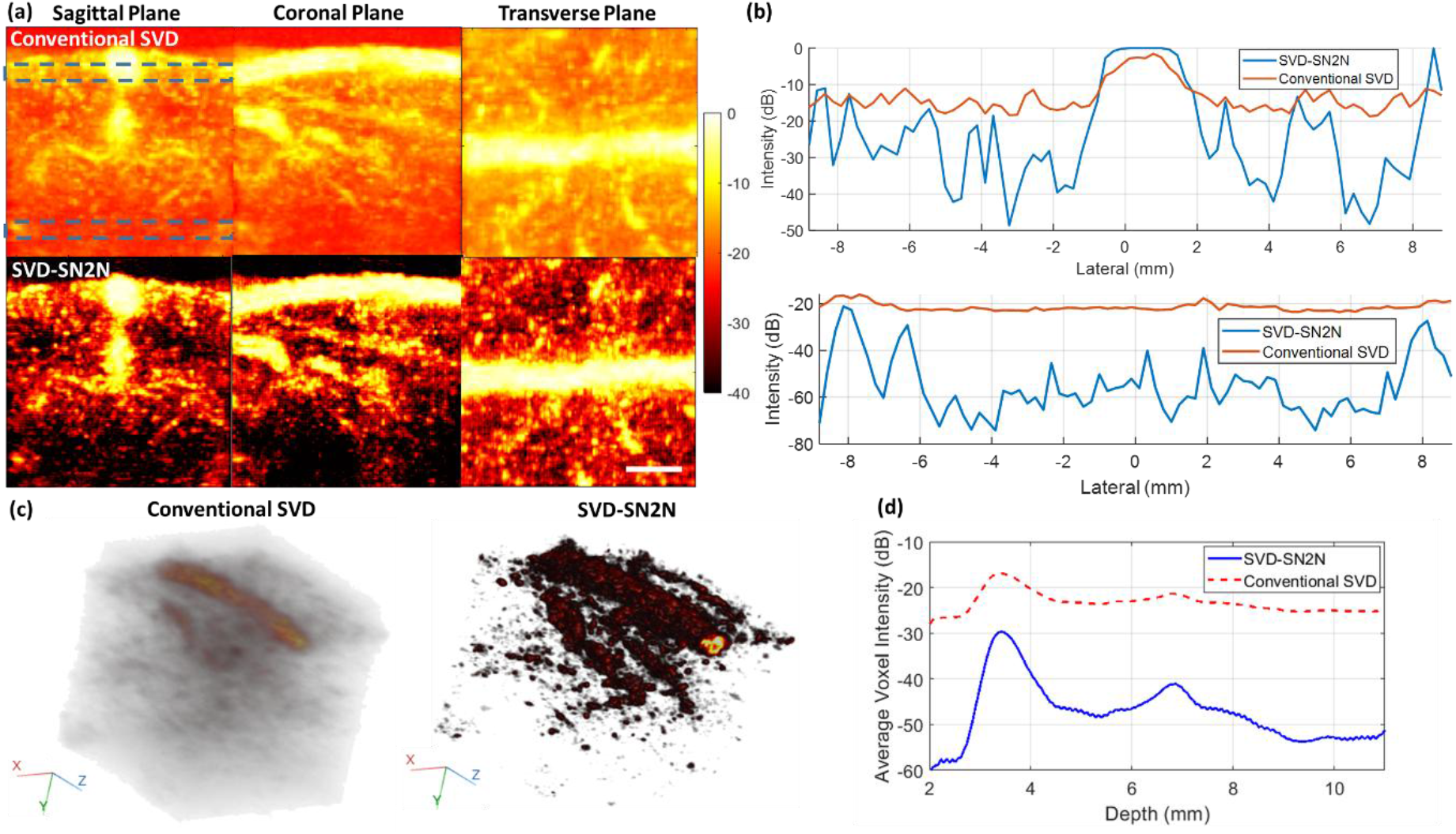
*In-vivo* 3-D rat-brain CEUS imaging acquired with a row-column-addressed array over 50 volumes (Dataset 4) at a mechanical index of 0.025. (a) Conventional SVD and log-compressed SVD-SN2N on the sagittal, coronal and transverse planes. (b) Intensity profiles along a lateral line through the near-field cortex region in the sagittal plane (top) and in the far field, 9–10 mm in depth (bottom). (c) Volume renderings for both methods. (d) Depth-resolved average voxel intensity. All images are displayed over a 40 dB dynamic range.

## 4. Discussion

The central methodological contribution of this work is the insertion of a log-compression step between the SVD filter and the self-supervised training. This step is not cosmetic: it changes the statistical nature of the residual noise from multiplicative speckle, which is signal-dependent and therefore incompatible with N2N theory [26], into an additive perturbation whose variance no longer depends on the signal. The diagnostics of Section 3.1 confirm this quantitatively (variance 0.409 against the predicted 0.411, equivalent looks 1.00) while making explicit that the residual is Fisher–Tippett rather than Gaussian, which the *ℓ*_1_ objective accommodates. The experimental behaviour follows the prediction: a network trained on linear-scale inputs yields outputs essentially as noisy as the input, whereas the same network trained on log-compressed inputs produces visibly cleaner images while preserving vascular structures.

Three numerical patterns deserve emphasis. First, the asymmetry between the two pipelines: with an identical network, loss, augmentation and optimizer, the log pipeline gains several decibels of SNR and narrows the apparent FWHM, while the linear pipeline gains at most 2.4 dB and does not narrow vessels at all. Since the only difference is the pre-processing, the asymmetry identifies log-compression, and not the network, as the operative ingredient. Second, the gain depends on the regime: it is small when the baseline SNR is already high and grows to roughly +10 dB on the lowest-SNR ROIs. This is what multiplicative-noise theory predicts, because the variance of a multiplicative perturbation scales with the signal amplitude and therefore dominates most severely where the signal is weakest. That this predicted dependence is reproduced across four ROIs spanning two imaging contexts and a baseline SNR range of 6–20 dB is the strongest empirical argument for the log-compression framing. Third, the spatial counterpart: in both datasets the deeper ROI is the lower-baseline-SNR ROI, because depth-dependent attenuation, aberration and out-of-focus speckle accumulate onto the residual noise floor. Log-compressed SVD-SN2N consequently lifts the deeper region more than the superficial one, and the superficial-to-deep gap shrinks from 6.4 dB to 0.4 dB on the rat brain and from 6.7 dB to 4.9 dB on the breast.

It is important to state precisely what these metrics do and do not establish. SNR and FWHM as used here are *image-derived*: they quantify the contrast between vessel and background regions of the displayed map, and the apparent width of structures in it. A narrower apparent FWHM is consistent with, but is not proof of, improved spatial resolution or faithful recovery of the underlying microvasculature, since a sufficiently aggressive denoiser can sharpen structures without adding information. We have not compared the denoised output against an independent structural reference such as a co-registered ULM reconstruction or histology, and we therefore make no claim of structural fidelity. The evidence presented here supports a narrower claim: that log-compression makes self-supervised denoising converge on SVD-filtered ultrasound images, and that the resulting maps have a lower background floor and better-delineated vessels than conventional SVD. Establishing that the recovered structures correspond to real vessels requires a reference-based validation that is left to future work. For the same reason the self-consistency map of Section 2.9 is reported as an uncalibrated relative indicator rather than as a validated uncertainty estimate.

To establish how broadly the framework generalizes, we tested it on four datasets differing in dimensionality (2-D vs. 3-D), in the presence and type of contrast agent, and in the underlying scanner (Verasonics, Iconeus One, Mindray Resona R9). The same behaviour is recovered in all four cases, indicating that the operative ingredient is the log-compression-induced linearization of the residual speckle rather than a property of one probe or scanner. A complementary practical advantage is that the method is designed around the data clinicians and researchers actually have: it operates entirely on the post-SVD image, treats the scanner-produced image as a noisy observation of the underlying log-compressed vascular signal, and requires no clean reference and only one to two seconds of acquisition. This makes it applicable to clinical data streams in which only beamformed or post-compression images are exposed.

### Clinical relevance

Three properties determine whether the framework can be used in practice, and each maps onto a concrete constraint of clinical ultrasound. First, it consumes only what a commercial scanner exports. The breast examination analysed here was acquired on a clinical system that gives no access to RF or IQ data, and the entire pipeline—SVD filtering, log-compression, self-supervised training and inference—was run on the exported DICOM contrast sequence alone. Deployment therefore requires no scanner modification, no research licence and no change to the acquisition protocol, and can be applied to examinations already recorded in routine practice, including retrospectively. Second, training is per-examination: two seconds of acquisition suffice, so no curated multi-patient dataset has to be assembled and no transfer between patients, probes or machines has to be assumed, since the model is refitted to the noise of the study being read. Third, the improvement is largest where reading is hardest. In the breast examination the image-derived SNR rose by 8.8 dB in the superficial tumour rim but by 10.6 dB in the deep region, and the superficial-to-deep gap narrowed from 6.7 dB to 4.9 dB, so that microvascular structures previously buried in the noise floor at depth became readable. Depth is precisely where contrast-enhanced assessment of lesion vascularity is least reliable and where lesions are most often left indeterminate.

These properties indicate where the method could plausibly matter—characterization of deep or strongly attenuating lesions, shorter contrast acquisitions, and cleaner inputs for microbubble localization—but they do not by themselves demonstrate clinical benefit, and we are careful not to present them as such. The clinical material analysed here is a single examination; no reader study was performed and no diagnostic endpoint was measured. Demonstrating clinical impact would require a multi-reader, multi-case study against a histopathological reference, in which the question is not whether the image looks cleaner but whether inter-reader agreement or diagnostic accuracy improves. We regard the present results as establishing the technical precondition for such a study rather than as evidence of clinical benefit.

Several design choices support these results. Applying the N2N concept downstream of the SVD filter and on the log-compressed image combines the removal of large-amplitude tissue signals with the linearization of the residual speckle, yielding a noise regime close to the one N2N assumes. The diagonal resampling exploits the oversampling of ultrafast Doppler images relative to the PSF, while the Fourier zero-padding interpolation preserves the band-limited nature of the imaging system without introducing pixel-to-pixel correlations. The self-consistency term plays a larger role here than in the original N2N framework because the effective training population is very small—in the extreme case a single frame—and it drives the solution toward a unique estimate.

SVD-SN2N has several limitations. Like any N2N pipeline it relies on the residual noise being zero-mean and pixel-wise independent; Section 3.1 shows that the residual carries a correlation of 0.39 between the two companions, so this condition holds only approximately, and strong spatially coherent artifacts—near-field reverberations, aberration-induced ghosts, acquisition-to-acquisition bias—are not treated as noise and may be preserved or enhanced. A model trained on one patient or probe is not guaranteed to transfer to another geometry; the practical remedy is to retrain on each new acquisition, which is inexpensive given the single-frame protocol. The framework currently operates on 2-D power-Doppler images, including 2-D slices extracted from 3-D volumes; a fully volumetric extension is straightforward in principle, since diagonal resampling and Fourier interpolation generalize to 3-D blocks. Finally, the quantitative validation rests on two datasets and a limited number of ROIs, and the clinical dataset consists of a single patient examination, so the reported figures should be read as an initial characterization rather than as a population-level performance estimate.

## 5. Conclusion

We have proposed SVD-SN2N, a self-supervised Noise2Noise framework for denoising SVD-filtered ultrafast ultrasound images. Log-compressing the SVD output turns the multiplicative speckle into an additive term of signal-independent variance, diagonal resampling with Fourier zero-padding interpolation produces two nearly independent copies of the same image, and a U-Net trained with a self-constrained N2N loss reduces the residual background while preserving vessel geometry. The method requires no clean reference, operates entirely at image level and can be trained from one to two seconds of acquisition. Across four heterogeneous datasets it raises image-derived SNR by 4.2–10.6 dB and narrows the apparent vessel FWHM by 34%–59% relative to conventional SVD, while a linear-scale control gains only 1.1–2.4 dB and does not narrow vessels. The advantage grows as the baseline SNR decreases, the signature of a multiplicative-noise regime and direct experimental confirmation that log-compression is the operative ingredient. Establishing that the recovered structures are faithful to the underlying microvasculature requires reference-based validation and remains future work.

## Ethics Statement

### Animal data (Datasets 3 and 4)

All animal procedures were performed in compliance with European Directive 2010/63/EU and the corresponding French national regulations, and were approved by the local animal ethics committee (Comité d’éthique en matière d’expérimentation animale No. 59, Paris Centre et Sud) under project authorization no. 25358-2020051019027581V2. Experiments were designed and reported in accordance with the ARRIVE guidelines.

### Public data (Dataset 1)

Dataset 1 is a publicly available benchmark released by Heiles *et al*. [7]; animal preparation, anaesthesia and the corresponding ethical approval are reported in the original publication. The submission metadata for this article declare the use of animal data.

### Human data (Dataset 2)

The clinical contrast-enhanced ultrasound examinations analysed in this work were from a previously reported study of breast masses [28], which were acquired at the Department of Medical Ultrasound, China Resources & Wisco General Hospital, Wuhan University of Science and Technology, Wuhan, China, between October 2021 and March 2022, as part of. The protocol was reviewed and approved by the Institutional Review Board of China Resources & Wisco General Hospital and was conducted in accordance with the Declaration of Helsinki. Written informed consent to participate, and to the publication of potentially identifiable images and data, was obtained from every patient.

## Acknowledgements

The authors would like to thank Dr. Huarong Ye, Department of Medical Ultrasound, China Resources & Wisco General Hospital, Wuhan, China, for help with the clinical acquisition.

## Appendix

This appendix reports the per-ROI measurements underlying Sections 3.3 and 3.4. For SNR, the same manually drawn vessel/background ROIs were used across the three compared methods; for FWHM, the same automatically extracted set of line profiles was used across the three compared methods.

## References

[1] M. Tanter and M. Fink, “Ultrafast imaging in biomedical ultrasound,” IEEE Trans. Ultrason., Ferroelectr., Freq. Control, vol. 61, no. 1, pp. 102–119, Jan. 2014.

[2] C. Errico et al., “Ultrafast ultrasound localization microscopy for deep super-resolution vascular imaging,” Nature, vol. 527, no. 7579, pp. 499–502, Nov. 2015.

[3] C. Demené et al., “Spatiotemporal clutter filtering of ultrafast ultrasound data highly increases Doppler and fUltrasound sensitivity,” IEEE Trans. Med. Imag., vol. 34, no. 11, pp. 2271–2285, Nov. 2015.

[4] E. Macé et al., “Functional ultrasound imaging of the brain: Theory and basic principles,” IEEE Trans. Ultrason., Ferroelectr., Freq. Control, vol. 60, no. 3, pp. 492–506, Mar. 2013.

[5] O. Couture et al., “Ultrasound localization microscopy and super-resolution: A state of the art,” IEEE Trans. Ultrason., Ferroelectr., Freq. Control, vol. 65, no. 8, pp. 1304–1320, Aug. 2018.

[6] J. Baranger et al., “Adaptive spatiotemporal SVD clutter filtering for ultrafast Doppler imaging using similarity of spatial singular vectors,” IEEE Trans. Med. Imag., vol. 37, no. 7, pp. 1574–1586, Jul. 2018.

[7] B. Heiles et al., “Performance benchmarking of microbubble-localization algorithms for ultrasound localization microscopy,” Nature Biomed. Eng., vol. 6, no. 5, pp. 605–616, May 2022.

[8] P. Song et al., “Improved super-resolution ultrasound microvessel imaging with spatiotemporal nonlocal means filtering and bipartite graph-based microbubble tracking,” IEEE Trans. Ultrason., Ferroelectr., Freq. Control, vol. 65, no. 2, pp. 149–167, Feb. 2018.

[9] K. G. Brown, D. Ghosh, and K. Hoyt, “Deep learning of spatiotemporal filtering for fast super-resolution ultrasound imaging,” IEEE Trans. Ultrason., Ferroelectr., Freq. Control, vol. 67, no. 9, pp. 1820–1829, Sep. 2020.

[10] L. Huang et al., “Self-supervised deep learning for denoising in ultrasound microvascular imaging,” Biomed. Signal Process. Control, vol. 122, 2026, Art. no. 110368.

[11] G. Zhang et al., “Amplitude-modulated singular value decomposition for ultrafast ultrasound imaging of gas vesicles,” IEEE Trans. Med. Imag., early access, 2025.

[12] G. Zhang et al., “Exploiting harmonic signature of gas vesicles in amplitude-modulated singular value decomposition for ultrafast ultrasound molecular imaging,” Phys. Med. Biol., 2026.

[13] B. Heiles et al., “Nonlinear sound-sheet microscopy: Imaging opaque organs at the capillary and cellular scale,” Science, vol. 388, no. 6742, 2025, Art. no. eads1325.

[14] A. Bertolo et al., “XDoppler: Cross-correlation of orthogonal apertures for 3-D blood flow imaging,” IEEE Trans. Med. Imag., vol. 40, no. 12, pp. 3358–3368, Dec. 2021.

[15] J. Hansen-Shearer et al., “Ultrafast 3-D ultrasound imaging using row-column array-specific frame-multiply-and-sum beamforming,” IEEE Trans. Ultrason., Ferroelectr., Freq. Control, vol. 69, no. 2, pp. 480–488, Feb. 2022.

[16] M. A. Lediju et al., “Short-lag spatial coherence of backscattered echoes: Imaging characteristics,” IEEE Trans. Ultrason., Ferroelectr., Freq. Control, vol. 58, no. 7, pp. 1377–1388, Jul. 2011.

[17] G. Matrone et al., “The delay multiply and sum beamforming algorithm in ultrasound B-mode medical imaging,” IEEE Trans. Med. Imag., vol. 34, no. 4, pp. 940–949, Apr. 2015.

[18] J. Lehtinen et al., “Noise2Noise: Learning image restoration without clean data,” in Proc. 35th Int. Conf. Mach. Learn., 2018, pp. 2965–2974.

[19] A. Krull, T.-O. Buchholz, and F. Jug, “Noise2Void—Learning denoising from single noisy images,” in Proc. IEEE/CVF Conf. Comput. Vis. Pattern Recognit., 2019, pp. 2124–2132.

[20] J. Batson and L. Royer, “Noise2Self: Blind denoising by self-supervision,” in Proc. 36th Int. Conf. Mach. Learn., 2019, pp. 524–533.

[21] X. Li et al., “Reinforcing neuron extraction and spike inference in calcium imaging using deep self-supervised denoising,” Nature Methods, vol. 18, no. 11, pp. 1395–1400, Nov. 2021.

[22] M. Weigert et al., “Content-aware image restoration: Pushing the limits of fluorescence microscopy,” Nature Methods, vol. 15, no. 12, pp. 1090–1097, Dec. 2018.

[23] L. Qu et al., “Self-inspired learning for denoising live-cell super-resolution microscopy,” Nature Methods, vol. 21, no. 10, pp. 1895–1908, Oct. 2024.

[24] O. Ronneberger, P. Fischer, and T. Brox, “U-Net: Convolutional networks for biomedical image segmentation,” in Proc. Med. Image Comput. Comput.-Assist. Interv., Cham, Switzerland: Springer, 2015, pp. 234–241.

[25] R. J. G. van Sloun, R. Cohen, and Y. C. Eldar, “Deep learning in ultrasound imaging,” Proc. IEEE, vol. 108, no. 1, pp. 11–29, Jan. 2020.

[26] O. V. Michailovich and A. Tannenbaum, “Despeckling of medical ultrasound images,” IEEE Trans. Ultrason., Ferroelectr., Freq. Control, vol. 53, no. 1, pp. 64–78, Jan. 2006.

[27] Y.-M. Lei et al., “Combined use of super-resolution ultrasound imaging and shear-wave elastography for differential diagnosis of breast masses,” Front. Oncol., vol. 14, 2024, Art. no. 1497140.

[28] G. Zhang et al., “Ultrasound super-resolution imaging for differential diagnosis of breast masses,” Front. Oncol., vol. 12, 2022, Art. no. 1049991.

[29] N. Zucker et al., “PhysiofUS: A tissue-motion based method for heart and breathing rate assessment in neurofunctional ultrasound imaging,” bioRxiv, 2024, doi: 10.1101/2024.09.22.614324.

[30] M. Vert et al., “Transcranial brain-wide functional ultrasound and ultrasound localization microscopy in mice using multi-array probes,” Sci. Rep., vol. 15, no. 1, 2025, Art. no. 12042.

[31] J. Lecoq et al., “Removing independent noise in systems neuroscience data using DeepInterpolation,” Nature Methods, vol. 18, no. 11, pp. 1401–1408, Nov. 2021.

